# Cortical parenchyma wall width (CPW) regulates root metabolic cost and maize performance under suboptimal water availability

**DOI:** 10.1101/2023.09.29.560009

**Authors:** Jagdeep Singh Sidhu, Ivan Lopez-Valdivia, Christopher F. Strock, Hannah M. Schneider, Jonathan P. Lynch

**Author notes:** **Author Contributions:** JSS, ILV, CFS, and HMS designed and conducted the experiments. JSS analyzed the data and wrote the manuscript. JSS and JPL conceived the idea. JPL obtained funding, supervised the project, and participated in experimental design and writing. **Competing Interest Statement:** Authors don’t have any competing interests.

## Abstract

We describe how increased root cortical parenchyma wall width (CPW) can improve tolerance to drought stress in maize by reducing the metabolic costs of soil exploration. Significant variation (1.0 to 5.0 µm) for CPW was observed within maize germplasm. The functional-structural model *RootSlice* predicts that increasing CPW from 2 to 4 µm is associated with *ca.* 15% reduction in root cortical cytoplasmic volume, respiration rate, and nitrogen content. Analysis of genotypes with contrasting CPW grown with and without water stress in the field confirms that increased CPW is correlated with ca. 32 to 42% decrease in root respiration. Under water stress in the field, increased CPW is correlated with 125% increased stomatal conductance, 325% increased leaf CO_2_ assimilation rate, 73 to 78% increased shoot biomass, and 92 to 108% increased grain yield. CPW was correlated with leaf mesophyll midrib parenchyma wall width, indicating pleiotropy. GWAS analysis identified candidate genes underlying CPW. *OpenSimRoot* modeling predicts that a reduction in root respiration due to increased CPW would also benefit maize growth under suboptimal nitrogen, which requires empirical testing. We propose CPW as a new phene that has utility under edaphic stress meriting further investigation.

**Significance Statement:** Suboptimal water availability is a primary constraint for global crop production that is intensifying due to climate change. The metabolic cost of soil exploration is a critical factor in plant performance under suboptimal water availability. This study highlights how increased root cortical parenchyma wall width (CPW) reduces root metabolic cost and improves crop adaptation to water deficit. Modeling results also indicate that increased CPW would be beneficial under suboptimal nitrogen availability. Therefore, CPW is a promising target for breeding crops with improved water and nitrogen use efficiency.

## Introduction

Suboptimal water and nitrogen availability are primary limitations to global crop production (1–4). During the 21^st^ century, almost every other year moderate to intensive drought covered more than 20% of agricultural land, annually affecting more than 1.5 billion people globally, leading to a multi-billion-dollar economic loss, food insecurity, and social turmoil (5–9). In low-input agroecosystems, nitrogen limitation is another major constraint to crop production, leading to increased risk of food insecurity and political instability (10–14). In high-input agroecosystems, intensive nitrogen fertilization is linked to significant economic and energy costs, as well as the deterioration of air and water resources (15, 16). Moreover, it is predicted that crop losses due to inadequate nitrogen and water availability will continue to rise in the future, especially in the face of increasing human population, climate change, and soil degradation (17). Developing crops and cropping systems with reduced requirements for water and nitrogen inputs is therefore a major goal for global food security (18–21). Screening for tolerance to suboptimal nitrogen and water availability is generally based on yield, however, yield-based selection has shown a low success rate because yield components are environmentally dependent and are mostly controlled by complex and unknown genetic pathways (22). Ideotype breeding employing assemblies of individual phene states (‘phene’ is to a ‘phenotype’ as ‘gene’ is to ‘genotype’, (23–25)) holds potential and has been proven effective to breed crop varieties with improved resistance to edaphic stress (22, 26–28). Notably, root phene states associated with reduced root metabolic cost improve plant performance under suboptimal water and nitrogen availability (29–32).

Root metabolic cost is a major constraint for efficient soil exploration under suboptimal water and nitrogen availability (32). Under these stresses, the root system can consume up to 50% of the daily photosynthate production for the construction and maintenance of living tissue as well as exudation (33–35). Plants also invest a significant portion (>25%) of scarce nutrients including nitrogen (N) and phosphorus (P) for root maintenance (35–37). It has been hypothesized that root phenotypes that reduce the metabolic costs of soil exploration can improve plant performance under drought and nutrient stress by enabling better capture of soil resources (4). Numerous empirical and *in silico* studies have corroborated this theory in important crops such as maize, wheat, barley, and common bean (as reviewed 10, 23, 28).

Root cortical aerenchyma consists of air-filled lacunae in the root cortex and is associated with reduced root respiration, nitrogen, and phosphorus content of root tissue, thereby reducing the metabolic costs of soil exploration and improving maize performance under drought, suboptimal nitrogen availability, and suboptimal phosphorus availability (10, 37–39). Similarly, reduced cortical cell file number and increased cortical cell size in maize are associated with reduced root respiration, nitrogen content, and phosphorus content which translates to improved growth under drought (10, 40–41) and suboptimal nitrogen availability (42). Mechanistically, these phenotypes decrease root metabolic cost by reducing the proportion of root tissue occupied by living cortical cells and active cytoplasm. Following this hypothesis, it has been proposed that subcellular phenotypes can also play an important role in regulating root metabolic cost and provide a novel suite of phenes for edaphic stress resistance (29, 43). Vacuolar and cell wall phenotypes are promising in this context since they occupy a substantial portion of tissue volume. An increased proportion of volume occupied by energy-efficient compartments including vacuoles and cell walls can potentially reduce root metabolic cost by reducing maintenance costs (4, 29, 44), which exceed tissue construction costs after a relatively short time (45).

Here we explore the role of cortical parenchyma wall width (CPW) in regulating root metabolic cost. Most plant cells with thick cell walls serve structural roles (46). Increased structural integrity is useful for plant adaptation to several abiotic stresses. Hypodermal and endodermal cell wall thickening and suberization are beneficial under salinity stress (47, 48), and increased cell wall width and lignin content in multiseriate cortical sclerenchyma (MCS) is beneficial under soil mechanical impedance in maize and wheat (49) and to some degree under drought in sorghum (50). These studies primarily focused on the width of cell walls in hypodermal or endodermal cells, leaving the functionality of cortical parenchyma cell wall width unexplored. Furthermore, the involvement of cortical cell wall width in regulating root metabolic cost has not been explored, which is the primary objective of this study.

Comparing different cell compartments, cell wall construction likely involves greater initial energy investment compared to cytosolic compartments, however, the maintenance cost of cell walls is relatively very low (51). Therefore, it’s reasonable to assume that an increased cell wall: lumen ratio would lead to reduced maintenance cost of a tissue. This mechanism of reduced tissue metabolic cost would provide a competitive advantage to a plant with increased CPW via reducing root metabolic cost and increasing soil exploration efficiency similar to reduced cortical cell file number and secondary root growth, and increased cortical cell size, root cortical aerenchyma and root cortical senescence (reviewed in 29). Furthermore, reduced root metabolic cost due to increased CPW would enable greater soil foraging and consequently be beneficial under edaphic stresses including drought stress and suboptimal availability of nitrogen and phosphorus. In this study, we assessed variation in CPW in maize germplasm and its adaptive value under drought stress in the field and under nitrogen limitation *in silico*.

For this study, we use maize as a model species. Maize is a leading global crop and is a staple food for approximately one-third of the human population, particularly in sub-Saharan Africa, Southeast Asia, and Latin America (52). Due to its widespread use as a staple food in various regions, as well as other factors, maize has become deeply ingrained in global agriculture, human diets, and cultural traditions (52, 53). Maize production is increasingly limited by suboptimal water and nitrogen availability (1, 54, 55), lending urgency to the development of maize cultivars with enhanced tolerance to suboptimal nitrogen and water availability. Furthermore, the availability of genetic resources in maize allows a better understanding and utilization of useful phenes like CPW. Additionally, the phylogenetic proximity of maize to other cereals like wheat, rice, and barley allows orthologous analysis and transferability of results (56).

The objective of this study was to test the hypothesis that increased CPW is associated with reduced root respiration, thereby leading to greater soil exploration, greater water capture, and improved resistance to drought. We first explore the value of increased CPW using functional-structural modeling in *RootSlice*. *In silico* predictions are then evaluated in the field. We also identify potential genetic loci linked with CPW. This manuscript represents a detailed analysis of the functional significance of CPW as a novel root anatomical phene in maize under drought stress. Additionally, *OpenSimRoot* (a functional-structural plant-soil model) predicts that CPW should also have utility under suboptimal nitrogen availability. The study aims to provide insights into the potential benefits of increased CPW for improving crop productivity under edaphic stress.

## Results

### Significant natural variation for CPW exists in maize

The Maize IBM (Intermated B73×Mo17) RIL population and the Wisconsin diversity panel were phenotyped to assess natural variation for CPW (Fig. 1A, B, C, and D). In both populations, CPW approximately follows a Gaussian distribution with a slight skew to the right (Fig. 1E and F). In the Wisconsin diversity panel, CPW varies by 3.8-fold (SD = 0.59 µm), ranging from 1.31 µm to 6.29 µm and a median of 2.54 µm (Fig. 1E). In the IBM RIL population, CPW varies by 2.5-fold (SD = 0.54 µm), ranging from 1.5 to 5.3 µm, with a median of 2.35 µm (Fig. 1F).

**Fig. 1.**
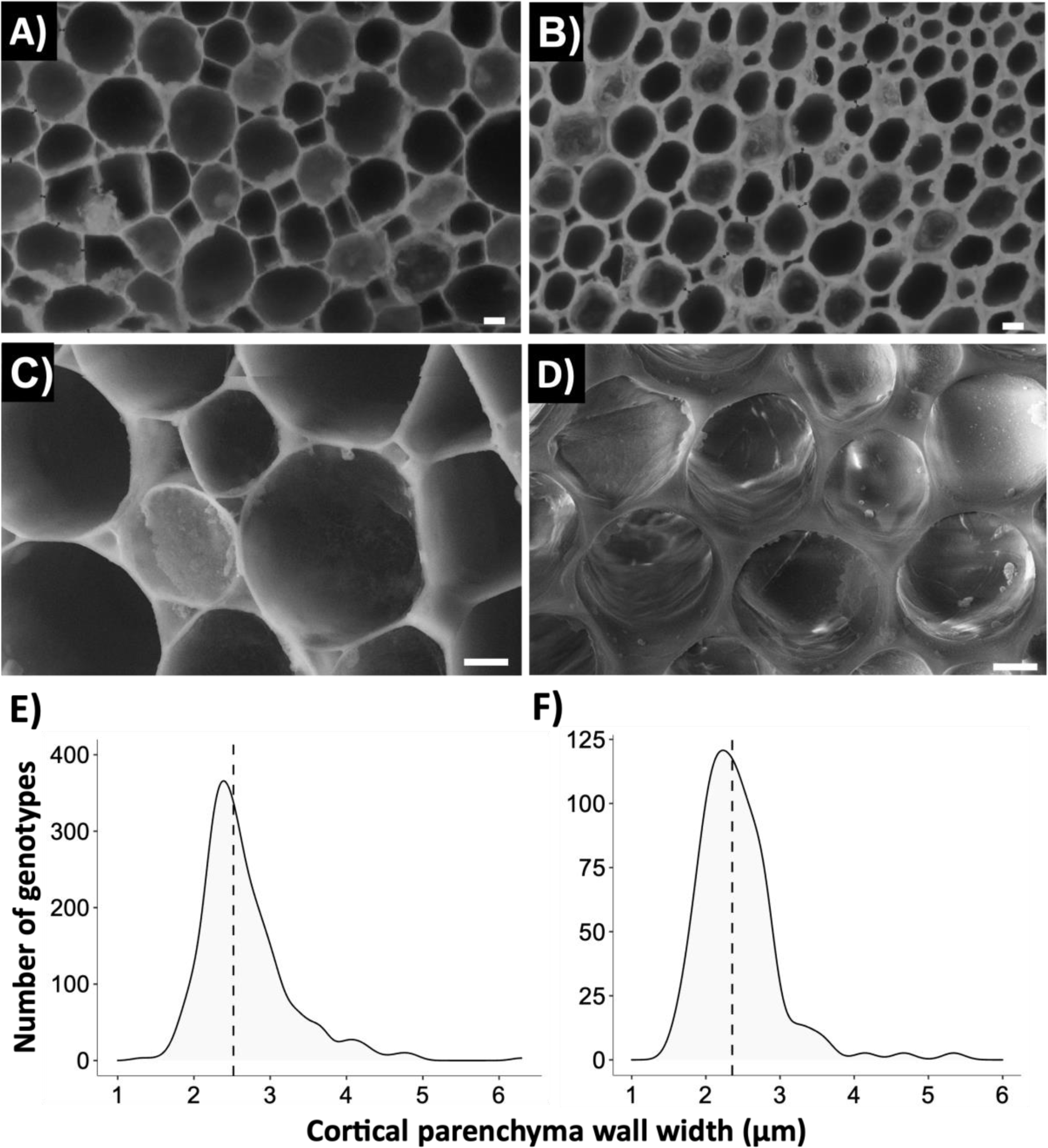
Natural variation for cortical parenchyma wall width (CPW) in maize germplasm. Laser ablation tomography images of a fourth node root sample representing thin CPW (A) and thick CPW (B). Cryo-SEM imaging of a thin CPW (C) and thick CPW (D) fourth node roots. Variation for CPW in the Wisconsin diversity panel (E) and the IBM-RIL population (F). Dashed lines indicate the median CPW for the respective population. Scales in A-D = 10 µm.

### *RootSlice* predicts that increased CPW is associated with reduced root respiration

Varying CPW from 1 µm to 4 µm was simulated using *RootSlice*-a functional-structural root anatomical model (Fig. 2). *RootSlice* predicts that with a 2 µm (from 2 to 4 µm) increase in CPW, the cortical volume occupied by cell wall content increases 176% (Fig. 2C) and cortical tissue density increases 84% (Fig. 2D). In contrast, the cortical volume occupied by cytosol declines 15% (Fig. 2E), thereby reducing respiration 15% (Fig. 2F), nitrogen content 17%, and phosphorus content 11.8% (Fig. 2G).

**Fig. 2.**
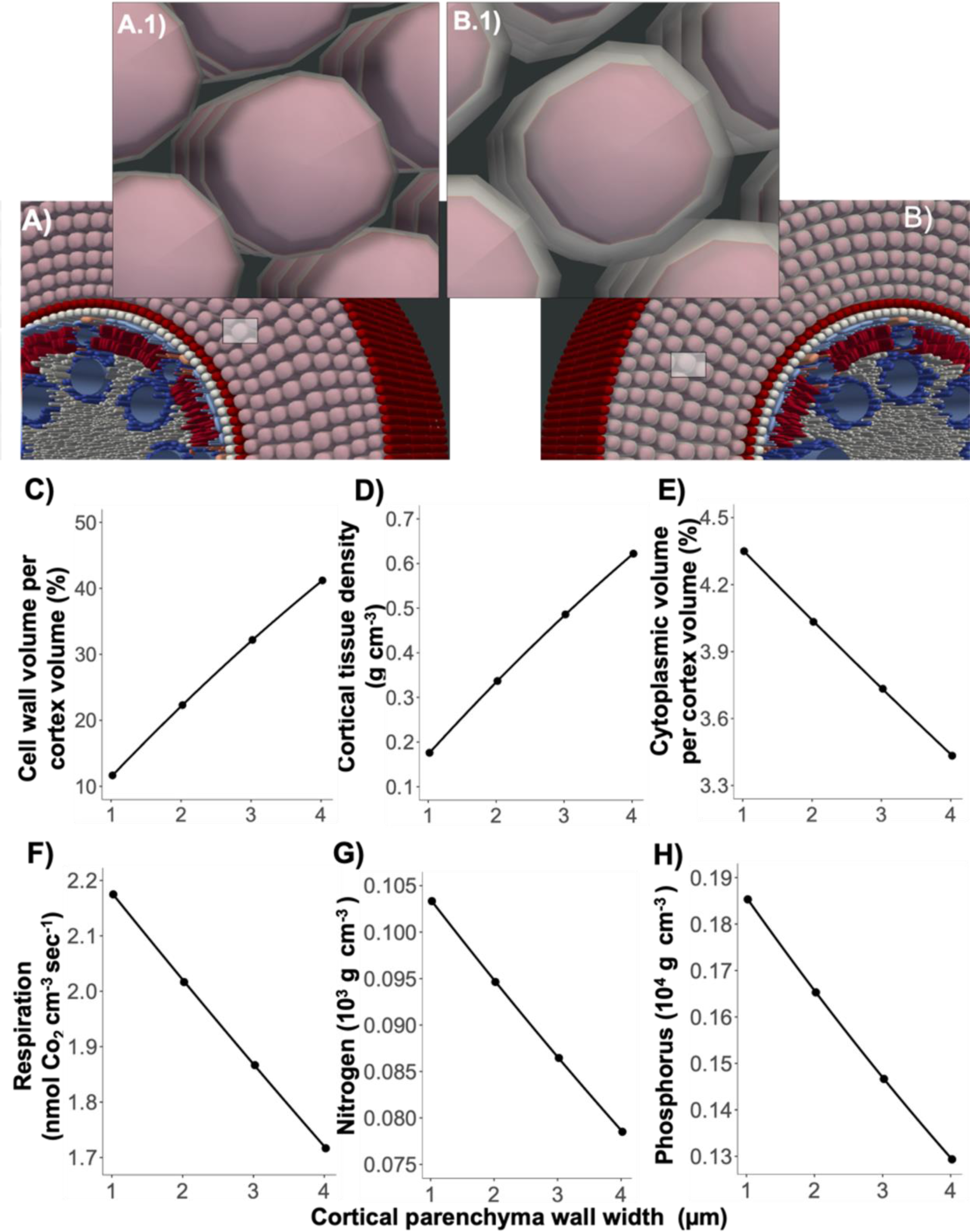
*RootSlice* shows the effect of cortical cell wall width (CPW) on tissue construction and metabolic cost. Maize root with 1 µm (A, A.1) CPW and 4 µm (B, B.1) CPW as simulated in *RootSlice*. Effect of CPW on root cortical cell wall volume (C), cortical density (D), cytoplasmic volume (E), respiration (F), nitrogen content (G), and phosphorus content (H). Note in Panel A and B, the grey color indicates cell wall and the pink color indicates cytosol.

### Increased CPW is associated with reduced respiration

A negative relationship between increased CPW and root respiration rate was found under both well-watered and drought conditions in the field (Fig. 3). Under well-watered conditions, CPW in this subset of IBM RIL genotypes ranged from 2.8 µm to 4.5 µm, and under drought it ranged from 2.4 µm to 4.7 µm. Increased CPW (2 µm, from 2 to 4 µm) under drought is associated with a 32% reduction in respiration per root dry weight (Fig. 3A) and a 42% reduction in respiration per root dry weight under well-watered conditions (Fig. 3B).

**Fig. 3.**
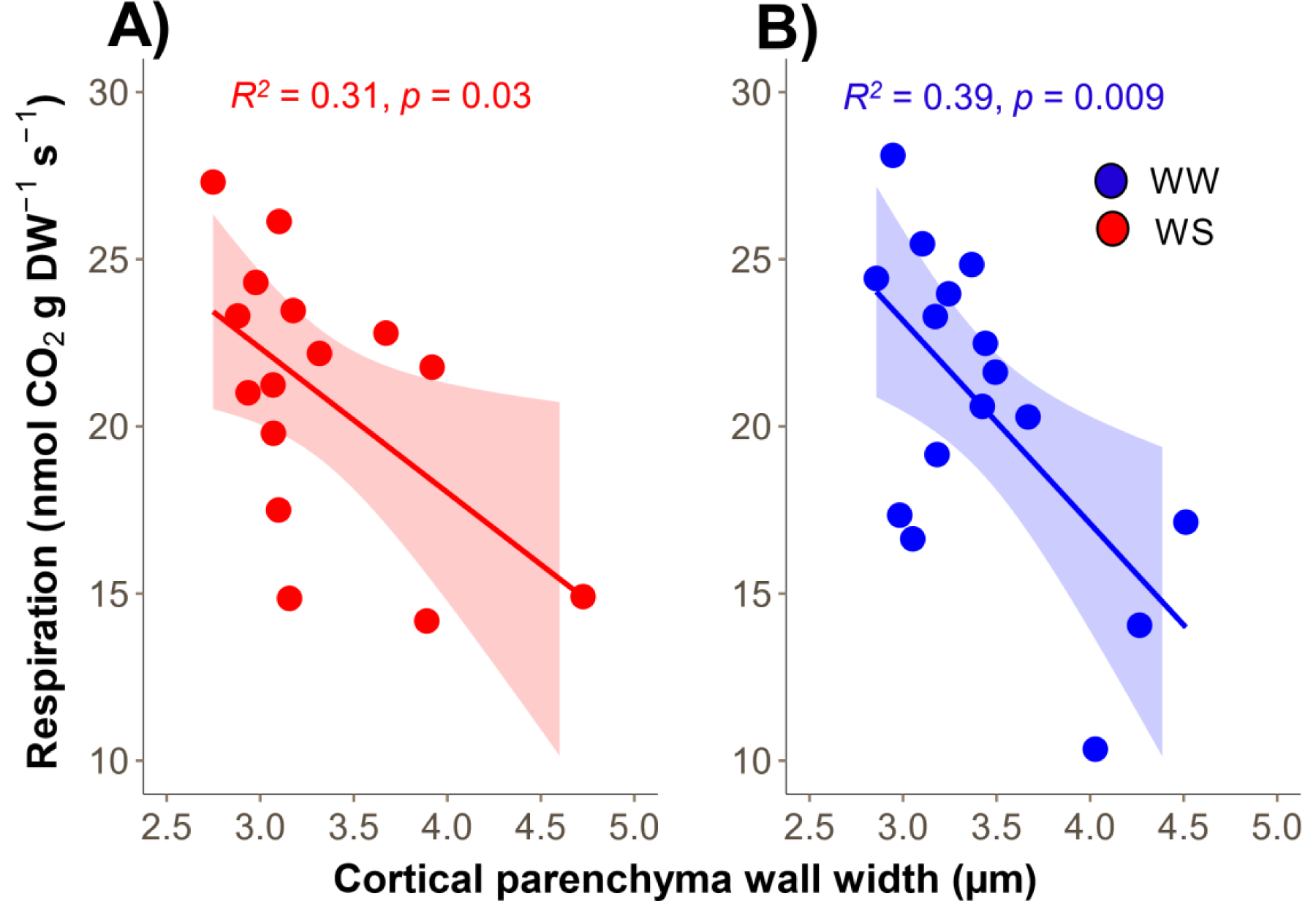
Relationship between cortical parenchyma wall width (CPW) and root respiration under well-watered (A) and water deficit conditions (B). Note that four IBM (intermated B73*Mo17) RILs (Supplementary Table S1) with varying CPW were grown in the greenhouse under two treatments (well-watered and water deficit conditions), each having four replications. Each point represents an individual measurement of a fourth node root from each replication (n = 15 for WS and 16 for WW). Lines represent best fitted linear regression line with color shading representing the 95% confidence interval around the mean.

### Increased CPW is associated with improved drought resistance in the field

The effect of varying CPW was tested under drought in the field in 2019 and 2021. In 2019 we used 14 IBM RIL lines with contrasting CPW phenotypes. In this set, under well-watered conditions, CPW ranged from 1.9 to 3.7 µm, and under drought CPW ranged from 1.7 to 4.0 µm. On average drought stress reduced shoot biomass by 37% and grain yield by 51% (Fig.4E, F, G, H). Under well-watered conditions, no significant relationship was found between CPW and response variables including transpiration rate (Fig. 4B, R^2^ = 0.12, p = 0.22), leaf carbon assimilation rate (Fig. 4D, R^2^ = 0.26, p = 0.06), shoot biomass (Fig. 4F, R^2^ = 0.03, p = 0.5) or yield (Fig. 4H, R^2^ = 0.0059, p = 0.4). Under drought, a positive correlation was found between CPW and growth metrics, where, increased CPW (from 2 to 4 µm) was associated with a 125% increase in mid-day transpiration rate (Fig. 4A, R^2^ = 0.39, p = 0.03), a 325% increase in carbon assimilation rate (Fig. 4C, R^2^ = 0.41, p = 0.03) a 73% increase in dry shoot biomass (Fig. 4E, R^2^ = 0.4, p = 0.01) and a 108% increase in grain yield under drought(Fig. 4G, R^2^ = 0.3, p = 0.05).

**Fig. 4.**
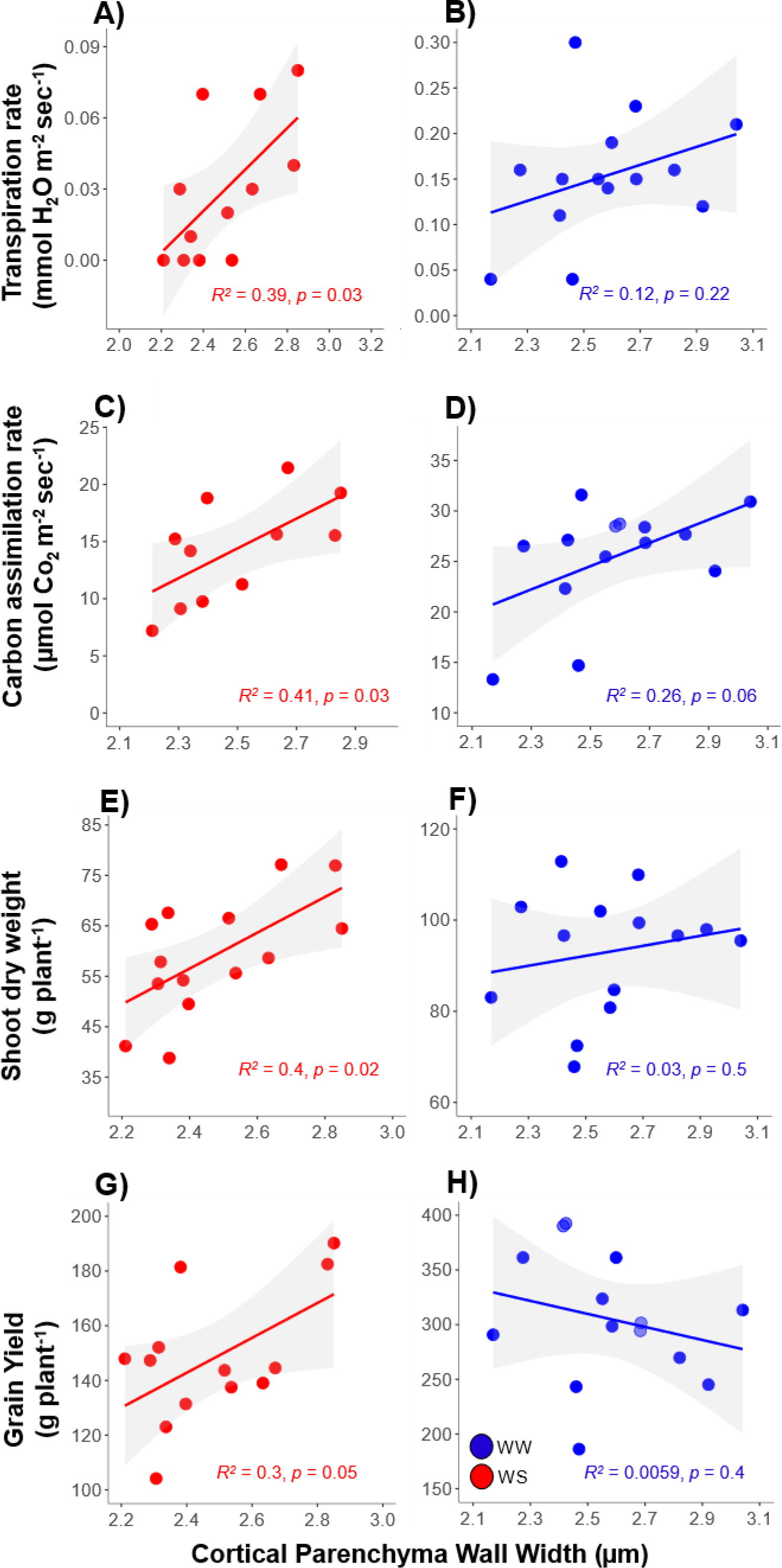
Effect of cortical parenchyma wall width (CPW) on plant performance in the field 2019 drought study. Transpiration rate (A, B), carbon assimilation (C, D), dry shoot biomass (E, F), and grain yield (include cobs, G, H) under well-watered and water deficit conditions, respectively. Each point is an average of four replications for each IBM (intermated B73*Mo17) RIL (n = 14). Lines represent best fitted linear regression lines with grey shading representing the 95% confidence interval around the mean.

In 2021, we used 7 IBM RIL lines with contrasting CPW phenotypes. In this set, CPW ranged from 1.74 to 5.82 µm and 1.68 to 4.83 µm under well-watered and water deficit conditions, respectively. Water deficit stress in this field study reduced shoot biomass production by 38% and grain yield by 34% (Fig. 5). Under well-watered conditions shoot biomass and CPW were unrelated (Fig. 5B, R^2^ = 0.091, p = 0.51) but under drought we again observed a positive correlation between shoot biomass and CPW (Fig. 5A, R^2^ = 0.59, p = 0.04). On average, a 2 µm increase in CPW leads to a 78% increase in shoot biomass. We also observed a positive relationship between CPW and yield under both well-watered (R^2^ = 0.68, p = 0.02) and water deficit conditions (R^2^ = 0.61, p = 0.04). To parse the effect of CPW on grain yield under drought, we estimated relative reduction in grain yield under drought compared to well-watered conditions. Interestingly, a negative relationship between CPW (R^2^ = 0.78, p = 0.0086) and relative reduction in grain yield under drought was observed (Fig. 5C). On average increased CPW (2 µm) is associated with 80% less reduction in grain yield under drought (Fig. 5C).

**Fig. 5.**
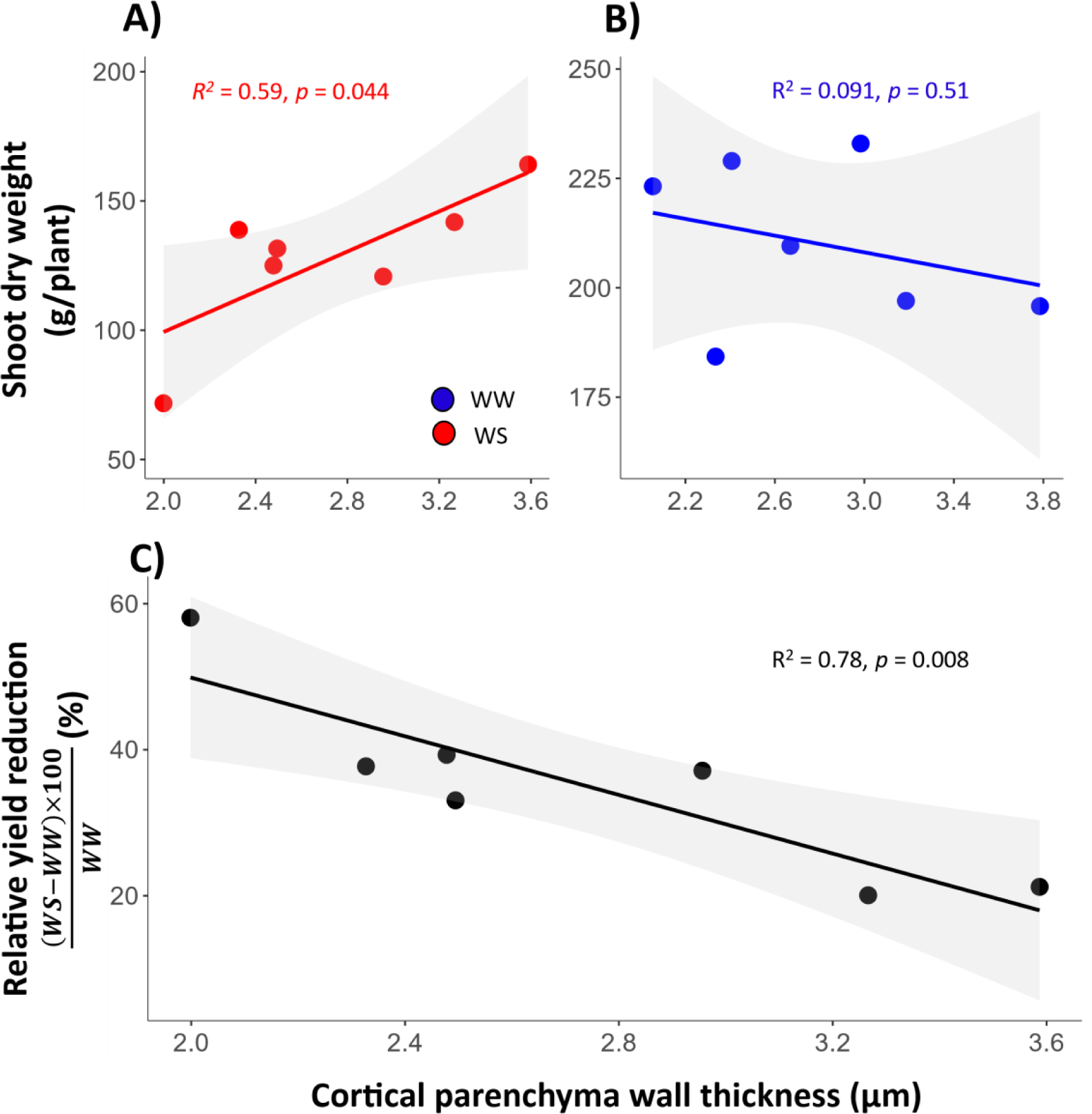
Effect of cortical parenchyma wall width (CPW) on shoot biomass and grain yield in the field 2021 drought study. Shoot dry weight under well-watered (A) and water deficit conditions (B). Relative reduction in grain yield due to drought stress (C). Each point is an average of four replications for each IBM (intermated B73*Mo17) RIL (n = 7). WS is water deficit and WW is well watered. Lines represent best fitted linear regression lines with grey shading representing the 95% confidence interval around the mean.

### Pleiotropy

The relationship between leaf midrib mesophyll parenchyma wall width (Fig. 6A) and root CPW (Fig. 6B) was assessed under well-watered and water deficit conditions. Leaf midrib mesophyll parenchyma wall width varied from 1.5 to 2.5 µm under drought and 1.4 to 2.6 µm under well-watered conditions. CPW ranged from 1.9 to 3.7 µm under well-watered conditions and 1.7 to 4.0 µm under drought. A positive linear relationship between root CPW and leaf midrib mesophyll parenchyma wall width was observed under well-watered conditions (Fig. 6D, R^2^ = 0.3, p = 0.04). In contrast, under drought, a weak relationship was observed between root CPW and leaf midrib mesophyll parenchyma wall width (Fig. 6C, R^2^ = 0.23, p = 0.08).

**Fig. 6.**
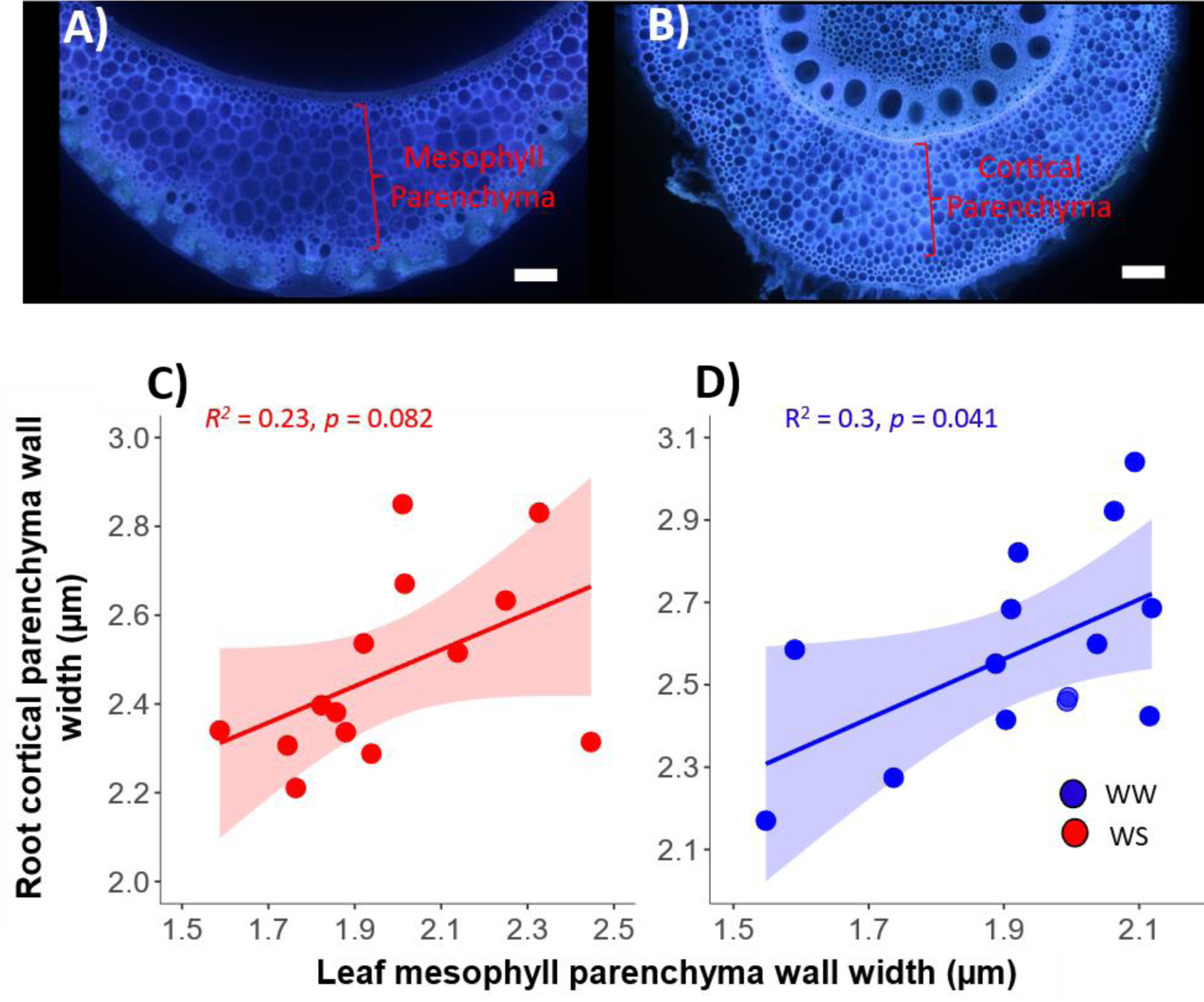
Relationship between root cortical parenchyma wall width (CPW) (A) and leaf midrib mesophyll parenchyma wall width (B) under water deficit (C) and well-watered conditions in the 2019 field experiment (D). An example of leaf midrib anatomy of the second youngest-expanded leaf (A), and 4^th^ node root anatomy (B). Each point is an average of four replications for each IBM (intermated B73*Mo17) RIL (n = 14). Lines represent best fitted linear regression lines with grey shading representing the 95% confidence interval around the mean. Scale in A and B = 100 µm.

### Genome-wide association mapping

CPW is heritable (H^2^ = 0.67) and under genetic control. A genome-wide association study (GWAS) using the BLINK method in GAPIT identified two significant SNPs (P < 0.000001) on chromosome 1 (Fig. 7A). The most significant SNP on chromosome 1 mapped to a D-3-phosphoglycerate dehydrogenase protein (*Zm00001d032114*). Under well-watered conditions *Zm00001d032114* is expressed in roots and leaves alike (www.maizegdb.org) and is highly expressed in the roots under drought (Fig. 7B). Another significant SNP on chromosome 1 mapped to a protein binding protein, related to the Ankyrin repeat family *(Zm00001d031706*). Under well-watered conditions *Zm00001d031706* is highly expressed in the roots, specifically in cortical tissue (www.maizegdb.org) and under drought it is highly expressed in root tissue (Fig. 7C).

**Fig. 7.**
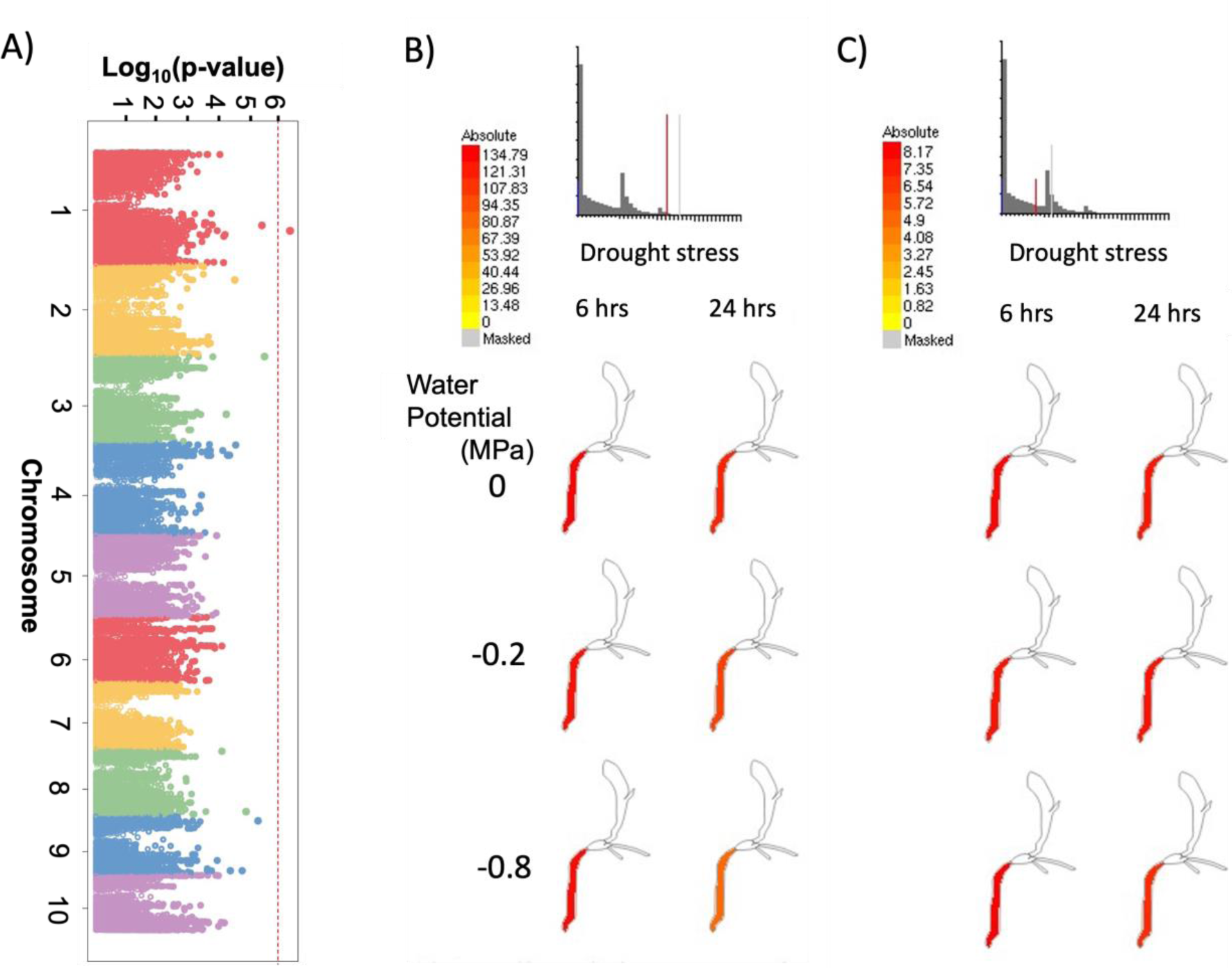
Genome wide association mapping for cortical parenchyma wall width (CPW). GWAS analysis results are represented as a Manhattan plot showing the−Log_10_(p) value of markers in the Wisconsin diversity panel (A). The expression profile of genes Zm00001d032114 (B) and Zm00001d031706 (C), which were found linked to the most significant marker on chromosome 1 under water deficit stress.

### *OpenSimRoot* predicts increased CPW improves resistance to suboptimal nitrogen availability

To understand the effect of CPW at an organismal scale we simulated thick (4 µm) and thin (2 µm) CPW phenotypes under varying nitrogen regimes using *OpenSimRoot* (Fig 8). Under nonstressed conditions (optimal water and nitrogen availability) we see no effect of CPW phenotypes on shoot growth. However, under suboptimal nitrogen availability, the thick CPW phenotype produced 106% more root length in soil domains deeper than 90 cm (Fig. 8D), resulting in 150% greater N capture (Supplementary Figure S4) and 87% greater shoot biomass compared to thin CPW (Figure 8C). Thin CPW phenotypes experienced stress 14 days earlier than thick CPW phenotypes (Figure 8E).

**Fig. 8.**
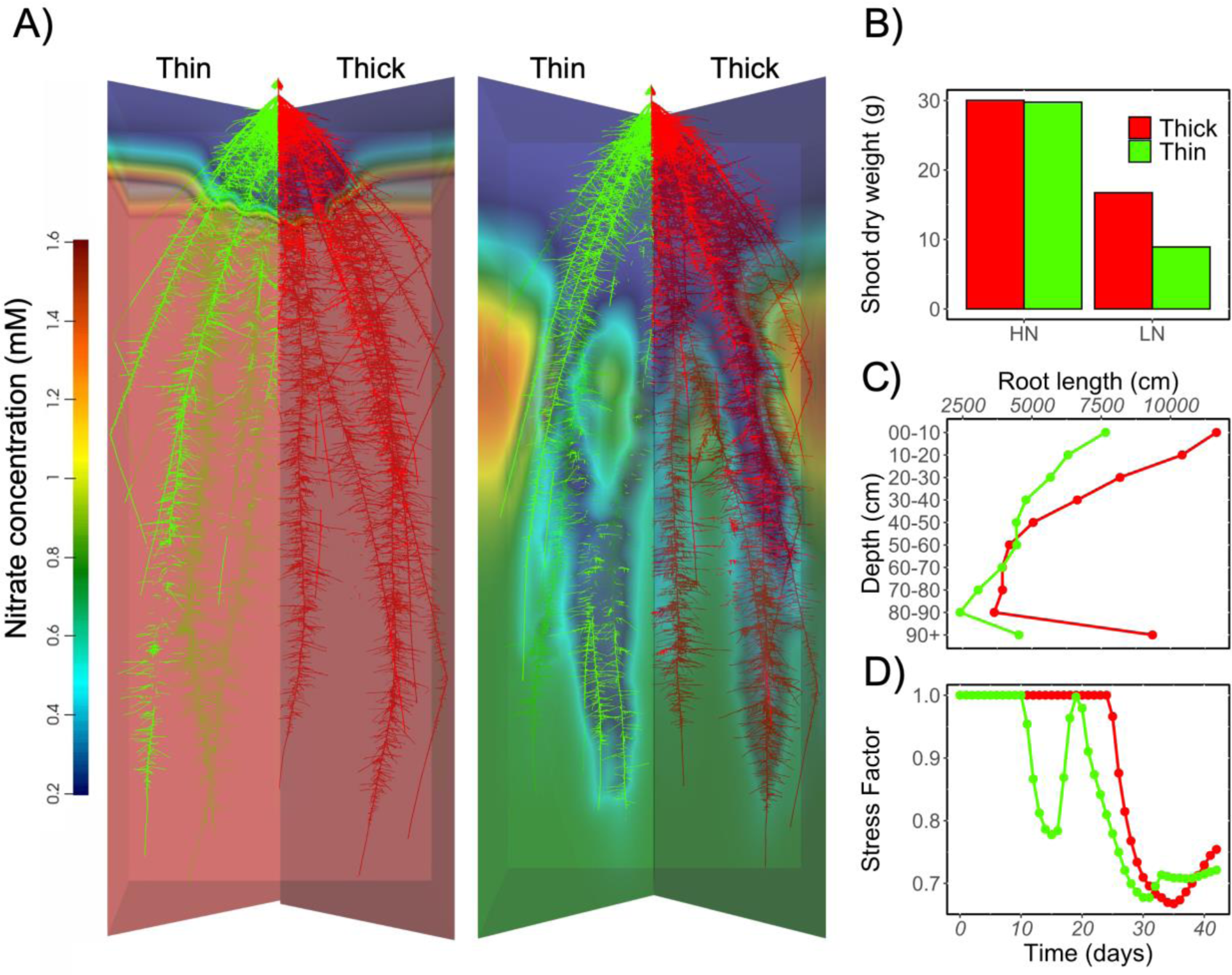
*OpenSimRoot* predicts increased cortical parenchyma wall width (CPW) is associated with improved plant performance under low nitrogen stress. Root system of thick (4 µm, red) and thin (2 µm, green) CPW phenotypes under optimum nitrogen and suboptimal nitrogen (A). Shoot dry weight (B), root length distribution (C), and stress factor profile (D) of thick (4 µm, red) and thin (2 µm, green) CPW phenotypes under suboptimal nitrogen availability.

## Discussion

Here we describe the utility of a novel root phene – cortical parenchyma wall width (CPW) – under suboptimal water and nitrogen availability. By screening a diverse set of maize genotypes, we show that significant natural variation exists for CPW in maize germplasm. *RootSlice,* a functional-structural model of root anatomy, predicts that increased CPW leads to reduced root respiration, which was empirically confirmed. Field experiments employing rainout shelters show that increased CPW is associated with greater transpiration rate, carbon assimilation rate, shoot biomass accumulation, and grain yield under drought stress. Under well-watered conditions, CPW was correlated with leaf midrib mesophyll parenchyma wall width, indicating pleiotropy. GWAS analysis reveals potential genetic loci and promising genes underlying CPW. *OpenSimRoot*, a functional-structural model of plant-soil interactions, predicts that decreased root respiration and root nitrogen content, as a result of CPW, would also improve nitrogen capture under nitrogen deficit.

The role of CPW in regulating root metabolic cost has not been explored. Root cortical cells are known to have varying cell wall properties based on their location in the cortex. Endodermal cell walls, for example, can be highly suberized and contain suberin lamellae which act as a barrier to passive transport into the stele (57). Similarly, hypodermal cells and sclerenchyma cells in the case of MCS have increased cell wall width and lignin deposition, which provides an adaptive advantage under soil mechanical impedance (49, 50). Helical cell wall thickenings of hypodermal tissue are reported in numerous plant species (58). However, the adaptive utility of variation in the cell wall width of cortical parenchyma cells has not been reported. Parenchyma cells are generally known to have thinner cell walls compared to sclerenchyma cells, however, variation in cell wall width can still be observed in parenchyma cells (59, 60). For example, leaf mesophyll parenchyma cell wall width varies among and between phylogenetic groups and influences photosynthesis (60). Drawing attention to CPW we here show that CPW influences root respiration and plant adaptation to suboptimal water and nitrogen availability.

Laser ablation tomography (LAT) (61) provides efficient high-throughput phenotyping of CPW. LAT measurements of CPW were confirmed by comparison with cryo-scanning electron micrographs (Supplementary Fig. S1), in accordance with a previous study (49). LAT measurements are mostly consistent with SEM images, with a slight overestimation of the CPW values which was corrected in this study based on the relationship between LAT and SEM phenotyping. However, some caution should be exercised in measuring CPW in samples with high phenolic compounds with strong autofluorescence (such as brace roots of maize, and nodal roots of sorghum and pearl millet species), which lead to low-resolution imaging and interference with CPW measurements in LAT. For these special cases, microtome sectioning followed by light microscopy (62, 63) or scanning electron microscopy provides a moderately efficient method to phenotype CPW. LAT is also highly efficient in obtaining leaf cross sections (64) and provides micron-level resolution to measure leaf parenchyma wall width. Leaf parenchyma wall width can be used to predict root CPW, however the relationship is influenced by environmental conditions. A positive relationship between CPW and leaf parenchyma wall width was observed under well-watered conditions (Fig. 6D) but a weaker relationship was observed under drought (Fig. 6C). Therefore, leaf parenchyma wall width may be used as a proxy measure for CPW, however, due to the context dependency of this relationship further confirmation is needed.

Significant variation (1 to 5 µm) for CPW was found within a maize inbred population and a maize diversity panel (Fig. 1). Similar trends have been observed in polypodiaceae with CPW ranging from 0.5 to 4.0 µm (58). However, much thinner (0.1 to 0.5 µm) cortical cell walls have been observed in *Arabidopsis* (65). Differences among species could be influenced by allometric scaling of cell wall width with cell size. Increase in cell size leads to increased turgor pressure (66) and larger cells may compensate for this pressure by developing thicker cell walls. However, larger cells are not always linked with thicker cell walls, in fact, an opposite relationship was observed in *Arabidopsis* where larger interfascicular fibers of an octoploid had thinner cell walls compared to smaller interfascicular fibers of a diploid (67). More research is needed to establish a clear link between cell size and cell wall width, as the relationship may vary depending on the species and cell type being studied. In maize, we show that differences in CPW are independent of cortical cell size (Supplementary Fig. S2). We also assessed the link between MCS and CPW. In the IBM and the Wisconsin diversity panel, we did not find a significant difference between MCS and non-MCS phenotypes for CPW (Supplementary Fig. S3C and S3D). However, we observed a positive correlation between sclerenchyma cell wall area: lumen area ratio and CPW under well-watered conditions in both field studies (Supplementary Fig. S3A and S3B) but under drought stress this relationship was either insignificant (p = 0.1, Supplementary Fig. S3B) or much weaker (R^2^ = 0.1, Supplementary Fig. S3A). Regardless of this relationship, only 7% of the samples analyzed had the MCS phenotype. Furthermore, MCS is observed only in the outer cortical layers but the CPW phenotype is present in the inner and middle cortex. Additionally, our genetic analysis reveals no common genetic loci between MCS (49) and CPW (Fig. 7), therefore we conclude that MCS and CPW are likely independent but related phenes.

Root metabolic cost plays an important role in regulating plant performance under edaphic stress including drought stress (32). Here, we show that CPW can regulate root metabolic cost. *RootSlice* predicted that an increase in CPW within the empirically observed phenotypic variation is associated with decreased root respiration and nitrogen content, and this reduction is due to an increased cell wall: cytosol volume ratio (Fig. 2C). We note the value of *in silico* biology in understanding novel phenotypes. *RootSlice* has played a central role in predicting and analyzing the value of CPW (44). *RootSlice’s* initial phenome space exploration and sensitivity analysis of CPW indicated the potential of CPW as a novel phene. *In silico* results were further validated both under well-watered conditions and drought, where an increase in CPW was associated with reduced root respiration (Fig. 3). Reduction in respiration due to increased CPW is analogous to the phenomenon of intracellular deposition of metabolically inert material leading to a lower metabolic rate (4, 43, 44, 68). For example, in animal cells metabolic efficiency can be achieved by the deposition of fat in adipose cells (69, 70). In plants, the accumulation of low-energy materials such as starch and water can similarly reduce tissue metabolic rate (71, 72). The involvement of energy-efficient vacuoles in larger cortical cells has been linked with reduced respiration and improvement in plant performance under drought and soil mechanical impedance (41, 73). The cell wall, once constructed, is also relatively metabolically efficient as compared to the energetically demanding cytosol and the organelles it contains (51). Therefore, increased width of parenchyma cell walls leads to reduced volume of the cytosol and in turn reduced root respiration. However, increased CPW leads to a substantial increase in tissue density (Fig. 2D), thus the construction cost of roots with thicker cortical cell walls would be greater. The carbon costs of maintaining a root tissue exceed the carbon cost of root tissue construction in a short growth period (33, 35). Furthermore, the maintenance costs can consume a significant portion of the daily carbon budget, especially under edaphic stress (36). Thus, the initial investment in increased CPW may pay off in the long term by reducing the daily root maintenance cost.

In the field, increased CPW was associated with better growth and yield under terminal drought stress (Fig. 4 and 5). Genotypes with increased CPW have greater leaf transpiration (Fig. 4A) and photosynthetic rates (Fig. 4C) at flowering, which translates to increased shoot biomass (Fig. 4E and 5A) and grain yield (Fig. 4G and 5C) under drought in two field studies performed in 2019 and 2021. In 2021, genotypes with increased CPW had greater grain yield under well-watered conditions as well, suggesting CPW may also be beneficial in high input agroecosystems. Increased CPW is linked to a reduced impact of drought stress on grain yield, i.e., less relative grain yield reduction indicating changes in CPW do not scale with allometry (Fig. 5C). Although it was not feasible to quantify rooting depth in this study, the maintenance of improved shoot water status and greater biomass is indicative of deeper rooting which allowed genotypes with increased CPW to acquire water from deeper soil domains under drought. Several other root anatomical phenotypes that reduce root metabolic costs including increased root cortical aerenchyma (38), increased cortical cell size (41) and reduced cortical cell file number (39) increase rooting depth and improve plant adaptation to drought. Furthermore, *OpenSimRoot* predicts that increased CPW is associated with deeper rooting under low nitrogen availability (Fig. 8D). We propose that increased CPW employs the same mechanism of reducing root metabolic cost to make roots metabolically cheaper and deeper. However, increased CPW might be relatively less beneficial under early drought stress as compared to terminal drought stress. The benefits of reduced respiration due to increased CPW are cumulative over time but might not outweigh the construction cost at earlier growth stages.

To test the utility of CPW under suboptimal nitrogen availability we employed *OpenSimRoot* (74, 75). *OpenSimRoot* is a functional-structural plant-soil model which considers tissue construction and maintenance costs and can simulate a wide variety of soil environments. *OpenSimRoot* has previously been employed to understand the utility of phenotypes that reduce root metabolic cost such as root cortical aerenchyma (37, 76), root cortical senescence (77, 79), larger cortical cell size (42), and reduced root cortical cell file number (42) under suboptimal nutrient availability and drought. *OpenSimRoot and RootSlice* are not predictive models but rather heuristic models testing the adequacy of a logic model and allowing a better understanding of the underlying mechanisms and functioning of root anatomical and architectural phenotypes based on the given parameters. In this study, both *RootSlice* and *OpenSimRoot* helped us explore the utility of CPW under resource-limited conditions. We simulated a range of CPW phenotypes with and without nitrogen stress (Fig. 8). Under optimum nitrogen (Fig. 8C), no differences in shoot biomass were observed among different CPW phenotypes. However, plants with thicker cortical cell walls outperform the plants with thin cortical cell walls under suboptimal nitrogen availability. (Fig. 8C). Thus, *OpenSimRoot* highlights that the benefits of reduced maintenance cost due to increased CPW outweigh the increased construction cost of cell walls. We further dissect the effects of individual functional components including root respiration, root tissue density, and root nitrogen construction cost *in silico* (Supplementary Fig. 4). Reduced nitrogen construction cost contributed 250% to the increase in root length, increasing nutrient uptake by 100%, and shoot biomass by 300%. Even though root respiration improved root length by 50% and plant nutrient uptake by 25%, there was no significant increase in shoot biomass. We observed that the thick CPW phenotype (root nitrogen construction + root respiration + root tissue density) improved root length by 300%, increasing nitrogen uptake by 140% and shoot biomass by 300%. The effect of the greater construction cost of thick CPW phenotypes is negligible, while the reduced nitrogen in tissue is the main factor contributing to the greater biomass compared with thin CPW. Reduced maintenance cost leads to deeper roots under suboptimal nitrogen (Fig. 8D). In most agroecosystems water and nitrogen accumulate in deeper soil domains, so increased CPW is potentially a valuable phenotype for many agroecosystems (4, 78). *OpenSimRoot* also shows that improved plant performance under low nitrogen stress is mainly driven by increased CPW and not by differences in shoot phenotypes. *OpenSimRoot* simulations employed isophenic models with varying CPW while the rest of the root and shoot phenotypes were kept constant. Therefore, the positive effects on plant performance under nitrogen stress can be directly attributed to increased CPW. Although predictions made by *OpenSimRoot* regarding the potential utility of increased CPW under suboptimal nitrogen availability were not tested in the field, the accurate prediction of numerous other related phenotypes (37, 77, 80) using *OpenSimRoot* highlights the potential for CPW to improve resistance to suboptimal nitrogen availability. Moreover, CPW belongs to a family (steep, cheap, and deep root system ideotype) of related phenes (4, 74) and it has been empirically shown that phenotypes from this ideotype are useful for both water and nitrogen deficit due to the mobile nature of these resources. Therefore, we propose that increased CPW merits attention as a strategy for improving plant performance under suboptimal nitrogen availability.

Existing variation for CPW within maize germplasm also hints at possible tradeoffs of increased CPW. Indeed, the construction cost of roots with thicker cortical cell walls is greater but this cost is outweighed by the benefits of reduced root respiration under drought. However, the interaction of CPW with other root anatomical and architectural phenes, such as root cortical aerenchyma or root loss, may alter the cost-benefit ratio, because these phenes can lead to cell death or root loss, which may affect the benefits gained from investing in thicker cell walls to reduce maintenance cost. Development of root cortical aerenchyma and root cortical senescence in roots with thicker CPW would reduce the benefits of increased CPW since the benefits of reduced maintenance cost could be dominated by loss of cortical tissue in roots with abundant root cortical aerenchyma and root cortical senescence. Similarly, the interaction of CPW with root loss deserves further investigation. Multiple studies highlight the impact of cell wall composition on crop resistance to biotic stress (81). We propose that cell wall thickness may also alter the susceptibility of roots to herbivores and pathogens by acting as a mechanical barrier to penetration. Root loss can occur due to abiotic stress, especially drought stress (82). Effects of root loss are context-dependent (83) and may vary for different edaphic conditions. The interaction of CPW with root cortical aerenchyma and root loss merits further exploration to identify optimum integrated phenotypes for specific stresses (40, 80, 84, 85). *In silico* tools like *RootSlice* and *OpenSimRoot* would be useful in exploring highly dimensional fitness landscapes as previously used to test the benefits of cortical cell file number and root cortical aerenchyma in maize (42–44).

The diversity in environmental conditions from which the analyzed genotypes were selected could also account for the observed variation in CPW across maize germplasm (86). Although the Wisconsin diversity panel consists of germplasm with restricted phenology, the panel includes genotypes from a range of environmental adaptations and has a significant representation of tropical alleles (86). Therefore, genotypes included have been bred and/or locally adapted to specific environmental conditions including soil taxa, nutrient availability, and water availability. Thicker CPW does not offer consistent benefits under optimum water and nutrient availability, thus explaining the prevalence of thinner CPW in maize germplasm. Nevertheless, variation in CPW can be utilized for selecting varieties with edaphic stress resistance. CPW in roots is associated with leaf mesophyll parenchyma wall width. However, the range of leaf mesophyll parenchyma wall width is much narrower and on average thinner than root CPW (Fig. 6). Increased leaf mesophyll parenchyma wall width might affect photosynthesis (60). It has been proposed that mesophyll CPW can limit mesophyll conductance (g_m_) – the ease of diffusion of CO_2_ from substomatal cavities to Rubisco active sites inside chloroplast stroma – which in turn can affect net CO_2_ assimilation. The potential mechanism by which cell wall width can reduce mesophyll conductance is based on the resistance to diffusion across the mesophyll cell wall. Cell wall conductance (g_cw_) is inversely related to the proportion of the cell wall resistance, which in turn is directly proportional to cell wall width. Therefore, along with other cell wall features such as porosity and tortuosity, cell wall width can add resistance to gas exchange. Other cell wall properties such as porosity and tortuosity are also at play and those properties are dynamic. For example, porosity can be changed by the reorganization of cellulose microfibrils and pectin deposition, but the cell wall width of the mesophyll cells might remain the same or change (60, 87). Furthermore, under drought we observed a weak non-significant relationship between CPW and leaf mesophyll parenchyma wall width which further highlights that increased CPW is advantageous under drought whereas increased mesophyll parenchyma wall width may not be. In a case where a positive relationship exists between CPW and leaf mesophyll parenchyma wall width under drought, the negative effect of thicker leaf mesophyll parenchyma walls could also be compensated by reduced root metabolic cost, as shown here. Under drought, we observed a positive relationship between CPW and photosynthetic rate (Fig. 4C). Thus, under drought, water capture is the major limitation to photosynthesis rather than mesophyll cell wall diffusion.

GWAS analysis reveals potential genetic loci underlying CPW (Fig. 7). Two significant SNPs on chromosome 1 were linked with the gene models encoding an Ankyrin repeat family protein and a D-3-phosphoglycerate dehydrogenase protein. Both genes are highly expressed in roots under drought stress (Fig. 7B and C). Even though both gene families are involved in cell morphogenesis, the Ankyrin repeat family protein acts upstream of cell wall organization or biogenesis (88). Furthermore, the Ankyrin-Repeat gene “*GmANK114*” confers drought resistance in *Arabidopsis* and soybean (89). Therefore, we propose two candidate gene models namely *Zm00001d032114* and *Zm00001d031706* on chromosome 1 of maize that might regulate CPW.

We anticipate that CPW might be beneficial for other edaphic stresses including suboptimal phosphorus availability. Root phenotypes that reduce the root metabolic cost of soil exploration are useful under multiple edaphic stresses, examples include increased root cortical aerenchyma (10, 31, 37, 76), increased cortical cell size (40–43), and reduced cortical cell file number (39, 42). Since CPW reduces root metabolic cost it is probable that increased CPW would be beneficial for phosphorus acquisition under suboptimal phosphorus availability. Structural-functional models such as *OpenSimRoot* and *RootSlice* will be helpful to explore the benefits of CPW under low phosphorus stress.

Increased CPW merits attention to improve the performance of important cereals like maize under edaphic stress. Increased CPW reduces the metabolic cost of soil exploration, thereby improving growth under drought. *In silico* modeling also predicts the utility of increased CPW under suboptimal nitrogen availability. CPW is heritable and under genetic control, and the identified genetic markers could potentially be utilized in breeding programs to select for increased CPW.

## Materials and Methods

### RootSlice parameterization

For simulating varying CPW phenotypes, we simulated a root with 8 cortical cell files, a stele diameter of 360 μm, an average cortical cell diameter of 8 μm, 0% cortical aerenchyma, and 8 metaxylem vessels with an average metaxylem vessel diameter of 20 μm. Different levels of CPW (1 to 4 μm) were simulated while keeping other parameters constant.

### *OpenSimRoot* parameterization

We used *OpenSimRoot* to simulate two contrasting phenotypes of CPW, thick CPW (4 µm) and thin CPW (2 µm). To parameterize the root construction and maintenance costs for each phenotype, we specified the root carbon density, nitrogen content and respiration. For the thick CPW phenotype, the carbon density, nitrogen content, and respiration were 0.16 g/cm³, 1470.7 µmol N g⁻¹, and 0.01 g CO₂ g⁻¹ DW d⁻¹, respectively. In contrast, for the thin CPW phenotype, these values were 0.09 g/cm³, 3822.5 µmol N g⁻¹, and 0.03 g CO₂ g⁻¹ DW d⁻¹, respectively. We simulated plant growth for 42 days in silt loam soil texture under medium nitrogen stress, as described in detail in Lopez-Valdivia et al. (2023) (43). The input files required to reproduce these simulations are available in the following GitHub repository: https://github.com/ilovaldivia/Sidhu_et_al_2023.git.

### Respiration measurements

The experiments for respiration measurements were conducted in a controlled greenhouse environment at University Park, Pennsylvania. Four genotypes (Supplementary Table S1) with varying CPW were grown in the greenhouse in a randomized complete block design involving two treatments (well-watered and drought), each having four replications. Mesocosm growth media consisted of (by volume) 50% commercial grade sand (US Silica), 25% vermiculite (Whittemore Companies Inc., Lawrence, MA, USA), and 25% sieved topsoil (Hagerstown silt loam topsoil, a fine, mixed, mesic Typic Hapludalf), air dried, crushed and sieved through a 4 mm mesh. 10 g of Osmocote plus fertilizer (Scotts-Sierra Horticultural Products Company, Marysville, Ohio, USA) consisting of (%): NO_3_^−^ (8) NH_4_^+^ (7), P (9), K (12), S (2.3), B (0.02) Cu (0.05), Fe (0.68), Mn (0.06), Mo (0.02), and Zn (0.05)) was mixed in growth media for each mesocosm (150 cm high, 15 cm in diameter). A plastic liner was installed in each mesocosm, and ca. 30L of growth media was poured into each column. Seeds (one per mesocosm) were directly planted at a depth of 2 cm and plants were grown for 40 days under a 16/8-h (light/dark) photoperiod, 40% relative humidity, and maximum/minimum temperatures of 28°C/23°C. Midday photosynthetic active radiation was approximately 900 to 1,000 μmol photons m^−2^ s^−1^. Natural light was supplemented from 06:00 to 22:00 with approximately 500 μmol photons m^−2^ s^−1^ from LED lamps. Both treatments were irrigated for up to 28 days and after that water was withheld in the drought mesocosms to induce terminal drought stress. Upon harvesting (40 days), the root system was washed and 2 cm long adventitious root samples from the 4^th^ node were suspended in 1 mL of nutrient solution and transferred to a liquid oxygen electrode chamber (Hansatech Instruments Ltd, United Kingdom). The rate of root respiration was measured over 5 min. After root respiration measurements, the root segment was dried and weighed. Anatomy samples adjacent to the root respiration positions were taken and stored in 75% ethanol. The anatomy samples were imaged using laser ablation tomography (61).

### Field studies

Field studies were conducted at the Russell E. Larson Agricultural Research Farm at Rock Springs, PA, USA (40°42’40.915” N, 77°,57’11.120” W) from June through September 2019 and 2021. A split-plot design with two irrigation levels (well-watered and drought) was employed. In 2019, two 0.02 ha rainout shelters were split into two 0.01 ha blocks, and two 0.02 ha irrigated fields were split into two 0.01 ha blocks. Fourteen IBM RIL genotypes (Supplementary Table S1) were randomized within each block, with 4 replications per treatment-genotype combination. Before planting, all fields were fertilized to meet the nutrient requirements of maize as determined by soil tests at the beginning of the season. Each genotype was planted in a 4.6 m long plot with 76 cm row spacing at a density of 73,300 plants ha^−1^. As needed, irrigation was supplied to the well-watered treatment. Drought treatment was initiated at 20 days after planting after which, drought plots experienced no rainfall or irrigation through the time of yield harvest. In case of rainfall events, drought plots were automatically covered. In 2021 we employed similar methods but used four rainout shelters instead of two, three-row plots instead of one-row plots, and 7 genotypes with contrasting CPW phenotype were used instead of 14 (Supplementary Table S1). At approx. 80 DAP (post anthesis), three plants per plot were harvested following the shovelomics protocol (90). Root anatomy samples include 4^th^ node roots (3 cm away from the base), fully opened youngest leaf lamina, and midrib. Anatomy samples were immediately stored in 75% ethanol and processed using laser ablation tomography (LAT) (61).

### Assessing variation in the IBM-RIL population

Variation for CPW was assessed in IBM (intermated B73 × M017) recombinant inbred lines (RILs) (91). The IBM panel was grown at the Ukulima Root Biology Center in Alma, Limpopo, South Africa (24.32002°S, 28.07427°E) from January to April in 2011. The experimental design was a randomized complete block design with four replications. In each rep, each genotype was grown in a single-row plot consisting of 20 plants per plot. The distance between plants within a row was 23 cm and the row width was 75 cm. Optimal growing conditions were maintained with additional irrigation applied using a center pivot system as needed. A subset of 137 genotypes was used for this study (Supplementary Table S1).

### GWAS for CPW

Genome-wide association study for CPW was performed using the Wisconsin diversity panel (86). The panel was originally grown at the Apache Root Biology Center in Wilcox, AZ, USA (32.1539252°N, 109.4956928°W) in 2016 under optimal growing conditions (40). At anthesis, one representative plant per plot was excavated and a 3 cm segment from 5 to 8 cm from the basal portion of the 4^th^ node root was excised from the crown for anatomical analysis. For this study, we re-phenotyped one replication of the fourth node root samples for CPW. The BLINK algorithm from the GAPIT R package was used to find marker-trait associations. Significant SNPs were identified based on a threshold of - log_10_(p-value) of 6. Significant SNPs identified in GWAS were translated to candidate genes based on the physical position of the genes in the version 4 B73 (AGPv4) reference sequence assembly (92). Expression analysis of the candidate genes was found in the MaizeGDB database (93).

### Cryoscanning Electron Microscopy

Cryo-scanning electron microscopy (cryo-SEM) was used to obtain images on a Zeiss Sigma VP-FESEM at the Pennsylvania State University Huck Institutes of the Life Sciences Microscopy Core Facility. Fourth node root samples were collected from the 80 day old plants grown in the field under drought stress. Root segments were stored in 75% ethanol in water (v/v) and then ablated and imaged using laser ablation tomography. On the day of cryo-SEM, each sample was mounted on a sample holder and plunged into liquid nitrogen. The sample holder was withdrawn under a vacuum into the cryo preparation chamber where the sample was maintained at −195 °C. The sample was then transferred to the SEM chamber onto a cold-stage module. Variable pressure without sputter coating was used. Voltage was 10 kV and samples were imaged at a temperature of −195 °C.

## Statistical analysis

The relationship between CPW and other response variables was assessed using linear models by employing the “lm” function from the “Stats” package in R (94). The significance of the linear models was determined at α = 0.05. CPW mean comparison among MCS and non-MCS phenotypes in the IBM and Wisconsin diversity panel was tested using the Wilcoxon test available in the R package “Stats” (94).

## Acknowledgments

We. acknowledge support from the US Department of Energy ARPA-E Award DE-AR0000821, the Howard G Buffett Foundation, and the US Department of Agriculture National Institute of Food and Agriculture and Hatch Appropriations Project PEN04732 and PENW-2020-03632.

## Data Availability statement

All the related data is available on Zenodo (DOI: 10.5281/zenodo.7767721)

