## Supplementary Data for "Cortical parenchyma wall width (CPW) regulates root metabolic cost and maize performance under suboptimal water availability"

This PDF file includes:

Supplementary Materials and Methods Figures S1 to S4

Tables S1

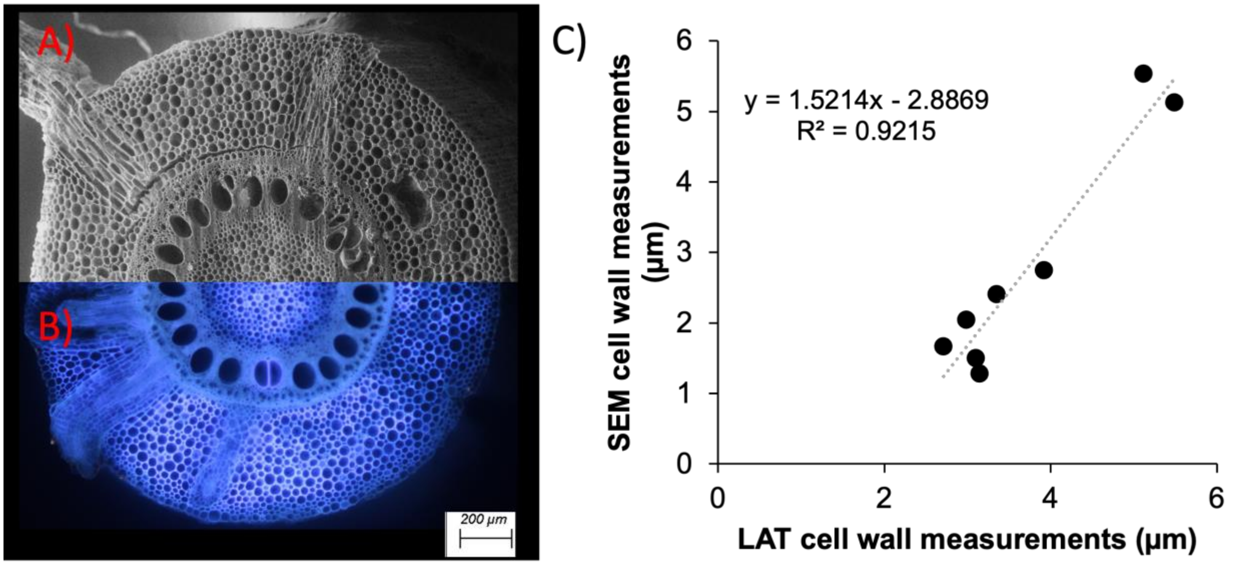


**Supplementary Fig. S1.** Validation of CPW measurements done using laser ablation tomography (LAT) with cryo-scanning electron microscopy (cryo-SEM). Root cross section imaged using LAT (A) compared with cryo-SEM micrograph (B). Relationship between LAT and cryo-SEM CPW measurements (C). Each point represents individual root sample first imaged using LAT and then imaged using cryo-SEM (n = 8).

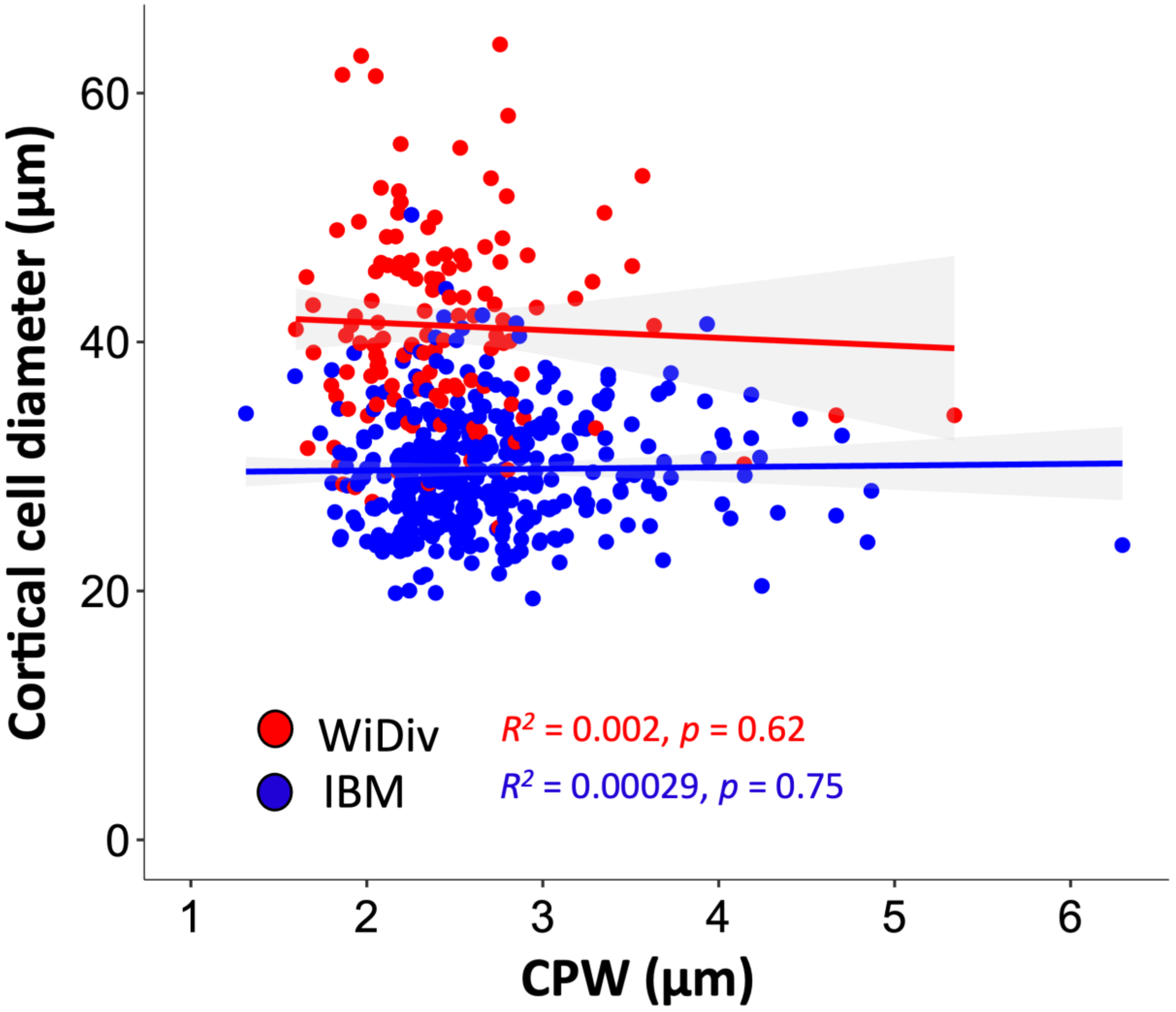


**Supplementary Fig. S2**. Relationship between CPW and cortical cell diameter in the WiDiv panel and the IBM population. Each point represents an average CPW and cortical cell diameter for each genotype (n for WiDiv = 365  n for IBM = 128s)

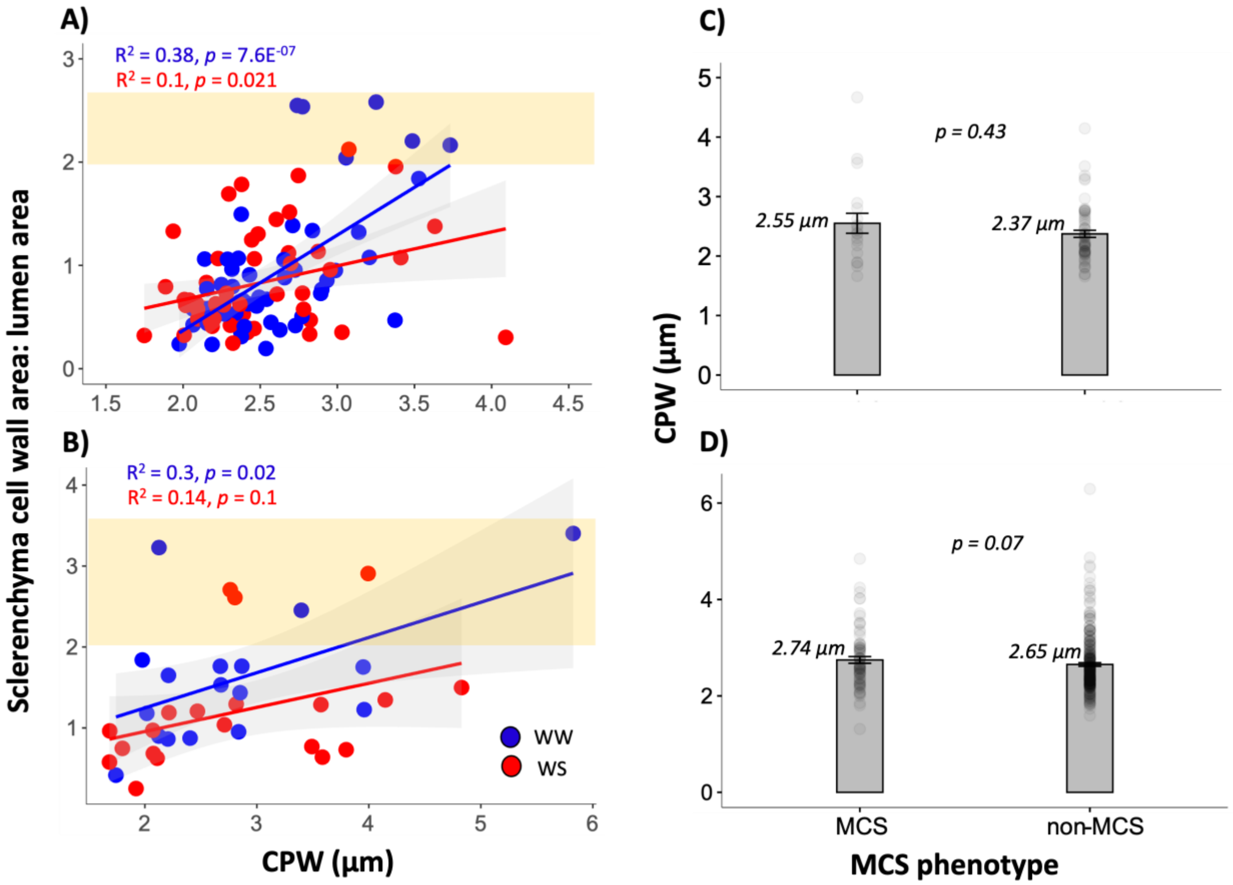


**Supplementary Fig. S3.** Relationship between CPW and sclerenchyma cell wall area: lumen area ratio under well water and water stress conditions in the 2019 (A) and 2021 drought study (B). Mean CPW comparison between MCS and non-MCS phenotypes in the IBM population (C) and the WiDiv population (D). Orange rectangles in A and B capture the points which classify as MCS.

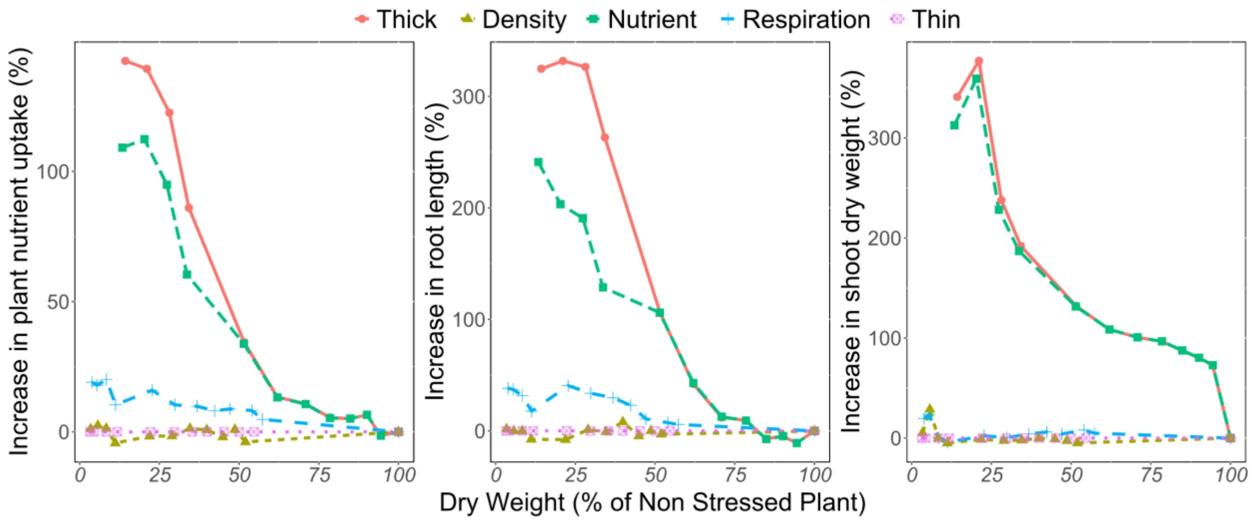


**Supplementary Fig. S4.** Relative effect of nitrogen construction cost (Nutrient), carbon construction cost (Density), and root respiration (Respiration). The x-axis represents the plant's performance under different levels of nitrogen stress, where 100% is the absence of stress. The y-axis represents the percentage of increase with respect to the thin phenotype. Thin and Thick phenotypes correspond to maize inbred lines with CPW of 2 µm and 4 µm, respectively.

**Supplementary Table S1.** Details about the genotypes used in the different experiments for this study. Note that IBM stands for Intermated B73*Mo17 RIL population and OHW stands for Oh43*W64A RIL population

| *Genotype* | *Experiment* | *Population* |
|---|---|---|
| *IBM001* | Drought 2019 | IBM |
| *IBM007* | Drought 2021 | IBM |
| *IBM010* | Drought 2019 | IBM |
| *IBM015* | Drought 2019 | IBM |
| *IBM017* | Drought 2021 | IBM |
| *IBM026* | Drought 2019 | IBM |
| *IBM111* | Drought 2019 | IBM |
| *IBM167* | Drought 2019 | IBM |
| *IBM177* | Drought 2019 | IBM |
| *IBM178* | Respiration | IBM |
| *IBM182* | Drought 2019 | IBM |
| *IBM200* | Drought 2021 | IBM |
| *IBM205* | Drought 2019 | IBM |
| *IBM284* | Respiration | IBM |
| *IBM284* | Drought 2021 | IBM |
| *IBM313* | Drought 2021 | IBM |
| *IBM317* | Drought 2021 | IBM |
| *IBM321* | Drought 2019 | IBM |
| *IBM327* | Drought 2019 | IBM |
| *IBM345* | Drought 2019 | IBM |
| *IBM351* | Drought 2019 | IBM |
| *IBM365* | Drought 2021 | IBM |
| *IBM379* | Drought 2019 | IBM |
| *IBM86* | Respiration | IBM |
| *OHW128* | Respiration | OHW |
